# Lipid flip flop regulates the shape of growing and dividing synthetic cells

**DOI:** 10.1101/2025.04.23.650179

**Authors:** Rafael B. Lira, Cees Dekker

## Abstract

Cells grow their boundaries by incorporating newly synthesized lipids into their membranes as well as through fusion of intracellular vesicles. As these processes yield trans-bilayer imbalances in lipid numbers, cells must actively flip lipid molecules across the bilayer to enable growth. Using giant and small unilamellar vesicles (GUVs and SUVs, respectively), we here recapitulate cellular growth and division under various conditions of transmembrane ‘flip flop’ of lipids. By dynamically monitoring the changes in reduced volume and spontaneous curvature of GUVs that grow by fusion of many small SUVs, the morphology of these growing ‘synthetic cells’ is quantified. We demonstrate that lipid flip flop relaxes curvature stresses and yields more symmetrically sized buds. Further increasing the neck curvature is shown to lead to bud scission. The mechanisms presented here offer fundamental insights into cell growth and division, which are important for understanding early protocells and designing synthetic cells that are able to grow and divide.

## Introduction

Living cells are characterized by a range of structural and functional elements that are fundamental to the cell’s ability to maintain homeostasis, reproduce, and evolve^1^. Paramount to the cell’s ability to proliferate is cellular division, a process by which a parental cell splits into daughter cells^2^. Morphologically, cell division is typically characterized by the progressive deformation of a cell at its midpoint, yielding a highly curved membrane structure that culminates in membrane scission and the generation of two daughter cells^2,3^ Although symmetric division is most common in cell biology, asymmetric forms of cell division also occur. For instance, in bacterial wall-less L-forms^4^, lipid overproduction yields an imbalance in surface area-to-volume ratio that drives cell shape deformations and lead to the extrusion of many small-sized L-forms, suggesting that growth and division can be driven by purely biophysical mechanisms without the need of a divisome protein machinery^5^.

In model lipid membranes of cell-like GUVs (giant unilamellar vesicles), the equilibrium shape of a vesicle is set by the interplay between the spontaneous curvature *m* of the lipid bilayer and the reduced volume *v = 6 V* √п */ A*^3/2^, where *V* and *A* are the vesicle volume and area, respectively^6–8^. This dimensionless parameter *v* describes how close a shape is to being to a perfect sphere, with *v* = 1 for a sphere. Notably, the vesicle equilibrium shape can be determined from an (*m*, *v*) shape diagram^9^ as it is controlled solely by biophysical parameters (*m*, *v*), independent of biological cues. How the two parameters *m* and *v* change during cellular growth depends on multiple factors. In most experiments using synthetic membranes, volume is regulated by osmotic changes, leading to changes in *v* at constant *m*^10^. In the case of growing membranes, lipid incorporation induces an increase in area (and accordingly a decrease in *v*), and in specific conditions, also to changes in *m.*^11^

Leaflet asymmetry is an important parameter of interest. For example, incorporation of lipid molecules into the inner leaflet of the bilayer membrane upon lipid synthesis in the cytoplasm^12,13^ leads to a change of *m* due to an imbalance in the lipid numbers across the membrane. Without a mechanism to balance this asymmetry, tension would build up due to an increased spontaneous curvature. As a result, cellular growth would stall as lipid incorporation would cause membrane budding rather than further expansion of the cell^14^. Common phospholipids exhibit only a slow transbilayer mobility, in the order of hours to days^15,16^, which is too slow to dissipate bilayer asymmetries during the timescale of biological processes such as cell growth. Therefore, cells have evolved ways to relieve transbilayer asymmetries. In fact, cells have flippases, enzymes that flip lipids across the membrane leaflets^17,18^, and it was recently demonstrated that cells can also dissipate bilayer asymmetries by spontaneously flipping cholesterol to the outer leaflet^19^. Such ‘flip flop’ mechanisms provide a way to quickly relieve membrane curvature and thus regulate cell shape.

Here we study these phenomena through an *in vitro* reconstitution approach. We examine whether rapid lipid flipflop is sufficient to alleviate bilayer stress and regulate the shape of growing and dividing synthetic cells. To study this, we fuse large amounts of small fusogenic SUVs (small unilamellar vesicles, ∼100 nm in diameter) to a GUV (>10 μm in diameter) to induce an asymmetry in lipid number across the GUV bilayer. Unlike GUVs, which are large and exhibit a nearly equal number of lipids in both leaflets, SUVs exhibit an intrinsic transbilayer asymmetry in lipid number, with an excess number of lipids in their outer leaflets^20^. In the absence of flip flop (i.e. when fusing SUVs to GUV membranes that only contain phospholipids), we observe GUV growth that is accompanied by a large decrease in *v*. As more and more SUVs fuse, a build-up in leaflet asymmetry yields a steep increase in *m* that gives rise to a budding transition where GUV growth is halted and the GUV instead forms a sphere that is decorated with many small outward buds. By contrast, when the GUV membranes contain a molecule that quickly undergoes flip flop, the gain in area is achieved with a much milder increase in *m*, resulting in the growth of more symmetrical GUVs with larger buds. By monitoring the fusion process in real-time, we quantify *v* and *m* as the GUVs grow, which enables dynamic tracking of the GUV shape trajectories in the *(m, v)* shape diagram in the absence and presence of lipid flip flop. These trajectories are found to be drastically different depending on whether lipids can flip. We furthermore show that inducing a further increase in *m* in budded vesicles can drive bud division. The results demonstrate that rapid lipid flip flop regulates membrane shape and alleviates curvature stresses caused by leaflet imbalances — which are crucial features for constructing synthetic cells from the bottom up that are capable of growth and division^21^.

## Results

### Fusion of SUVs deforms and buds GUVs

We tested the hypothesis that rapid lipid flip flop can alleviate transbilayer lipid-number asymmetries and regulate vesicle shape in growing and dividing synthetic cells. Our approach is to fuse SUVs, which exhibit a lipid asymmetry across both leaflets, to GUVs, which have a symmetrical lipid distribution. Fusion of SUVs with GUVs should induce a transbilayer lipid asymmetry in the GUVs, with an extent that scales with the number of fused SUVs (Figure 1A). Before SUVs addition, the GUVs are non-strained and spherical (Figure 1B, state i). In the absence of flip flop, fusion should lead to an increase in area and the appearance of membrane fluctuations, generating larger and non-spherical GUVs (Figure 1B, states ii and iii)^6–8^. Above a certain threshold, lipid asymmetries generated by SUV fusion should result in a budding transition, which is marked by a change from a floppy and non-spherical GUV to a spherical and tense GUV with outward buds. The initial buds are expected to be relatively large, but to become progressively smaller as more SUVs fuse (states iv and v in Figure 1B). We hypothesize that, for a given amount of SUV fusion, the presence of a membrane molecule that is able to quickly undergo flip flop and populate the lower-density inner GUV leaflet, would favour the formation of non-strained GUVs with smaller curvature effects, which could be assessed from their remodelling (i.e. shape, bud size).

**Figure 1.**
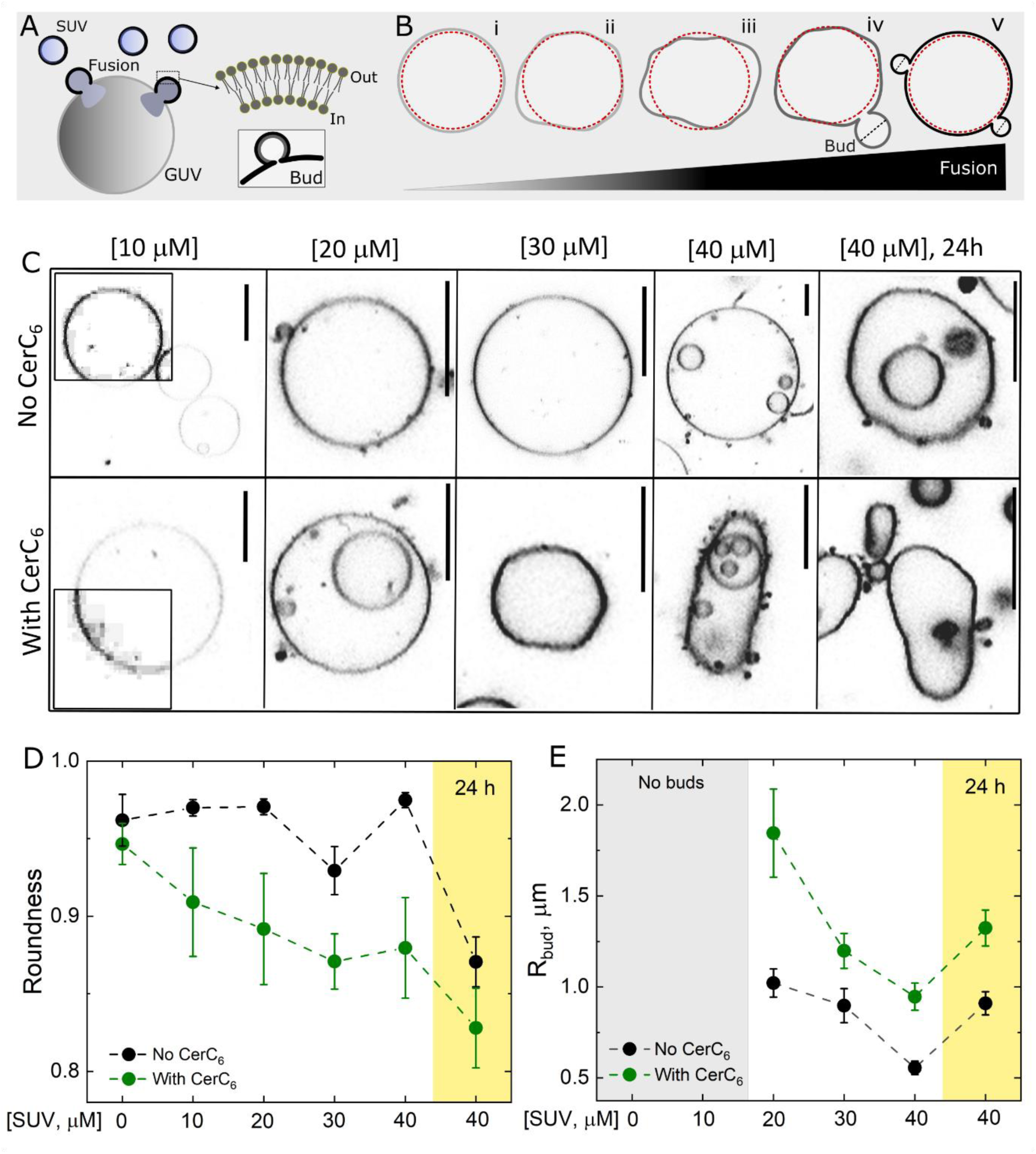
Lipid flip flop modulates vesicle shape and curvature. A, sketch of the SUV-GUV fusion experiment. An imbalance in lipid number across the outer (Out) and inner (In) leaflets of the SUVs is shown. Inset: a GUV bud formed after fusion. B, expected GUV shapes as fusion increases upon increase in [SUV]. Dashed red circles represent a spherical object, overlayed to progressively non-spherical GUVs. Note that the bud size decreases for the largest [SUV]. C, representative fluorescence images from GUVs devoid of (upper panel) or containing (bottom panel) CerC_6_ as a function of [SUV]. Insets at [10 μM] show contrast-enhanced images. Scale bars 20 μm. D and E, measured GUV roundness and bud radius R_bud_ as a function of [SUV] for GUVs devoid (black) or containing (green) 10 mol% CerC_6_. Yellow regions: GUVs observed 24h hours after SUV incubation to allow for slow and spontaneous lipid flip flop. Means and s.e.m are shown.

We set up an assay to test these scenarios. We fused highly fusogenic cationic SUVs of ∼60 nm radius^22^ made of DOTAP:DOPE (1:1 mol ratio), with negatively charged GUVs with a typical diameter of 10-40 μm made of DOPC:DOPG (1:1 mol ratio). The SUVs and GUVs were made by extrusion and electroformation, respectively (Methods). Assuming a bilayer thickness of 4 nm, the SUVs are expected to contain 14% more lipids in their outer leaflets, i.e. bear a lipid asymmetry of ∼1.14 across both leaflets. Fusion of SUVs to GUVs was assessed by the transfer of the fluorescent marker present in the SUV bilayers to the GUVs. As a model ‘flipping molecule’, we used CerC_6_, which was reconstituted in both the GUVs and SUVs at 10 mol%. The SUVs were made of DOTAP:DOPE:CerC_6_ (5:4:1 mol ratio) and the GUVs were made of DOPC:DOPG:CerC_6_ (4:5:1 mol ratio). We chose CerC_6_ for its ability to quickly (millisecond timescale) undergo spontaneous flip flop without enzymatic assistance^23^, which thus can be expected to relax leaflet asymmetries and regulate vesicle shape^24,25^. In principle, other molecules such as cholesterol could be used as well. However, we chose CerC_6_ over cholesterol because of cholesterol’s known effects on modifying the membrane mechanics^26,27^.

Figure 1C shows typical GUV morphologies as a function of incubated SUV concentration [SUV] for membranes containing or not containing CerC_6_. As [SUV] increased and more fusions occurred, the GUVs became less spherical due to the gained membrane area. Interestingly, we observed that the CerC_6_-containing GUVs were more floppy (yielding non-spherical GUVs) than the CerC_6_-devoid GUVs. We quantified the GUV roundness (see methods), i.e. a measure of how closely the GUV shape is to a perfect sphere, for GUVs containing or not containing CerC_6_ and for increasing [SUV] concentrations. Figure 1D shows that most CerC_6_-devoid GUVs were found to be near-perfectly spherical (roundness >0.94) for all [SUV] concentrations. For the CerC_6_- containing GUVs, however, significantly lower values were observed for the roundness – confirming the hypothesis that flipflop of lipids relaxes membrane tension. While most GUV shapes were evaluated after 10-15 minutes incubation with SUVs, Figure 1D also shows typical images (yellow shaded regions) for GUVs that were similarly incubated with the highest SUV concentration for 10-15 minutes, after which the SUVs were removed and the GUVs were allowed to relax for 24h (i.e. enough time for spontaneous slow flip flop). Here, a large GUV fraction was found to be non-spherical, irrespective of the inclusion or exclusion of CerC_6_.

Another morphological feature of curvature is budding. GUVs with a high transbilayer lipid-number asymmetry do bud as a result of the spontaneous curvature *m*, with *m = 1/R_bud_*, where *R_bud_* is the bud radius^28^. Thus, as more SUVs fuse, one would expect that the spontaneous curvature of the GUV increases and buds become smaller. This trend was indeed observed, both for membranes containing or devoid of CerC_6_ (Figure 1E). Importantly, the buds formed in GUVs containing CerC_6_ were always significantly wider than those in GUVs devoid of CerC_6_, again indicating that CerC_6_ alleviates curvature effects. When GUVs were allowed to relax for 24 hours after SUV removal, *R_bud_* increased. The results demonstrate that flip flop, even in the absence of CerC_6_, effectively alleviates leaflet stress. Both the effects of a flipping lipid on short timescales and slow flip-flop in the absence of CerC_6_ on long timescales reduce asymmetry stress, which leads us to conclude that lipid flip flop governs the shape of growing GUVs by alleviating transbilayer asymmetries and favouring less tense (more floppy) GUV shapes with larger formed buds.

### Shape trajectories of growing GUVs

Next, we followed fusion in real-time to dynamically track GUV shape changes as fusion progressed (i.e. more SUVs were fusing). Fusion of SUVs labelled with DOPE-Rh (yellow) with Atto- 390-labelled GUVs (blue) resulted in lipid mixing, as sketched in Figure 2A. The change in fluorescence intensity on the GUV was used to assess the number of SUVs (nSUVs) fused to a single GUV^29^. These experiments were similar to those from Figure 1 but the process now was observed in real time and the GUV history was known, which are necessary to assess *v* and *m*. Figure 2B shows examples of lipid mixing and changes in vesicle morphology versus time for GUV membranes devoid of or containing CerC_6_. The respective videos are shown in movies S1 and S2. As previously reported^30^, SUV fusion resulted in an increase in GUV area, the appearance of clear membrane fluctuations, and the GUV attaining non-spherical shapes. As more SUVs fused, the GUV underwent a budding transition. As even more SUVs fused, however, the GUV became spherical again, and the gained area was stored in buds that progressively decreased in size.

**Figure 2.**
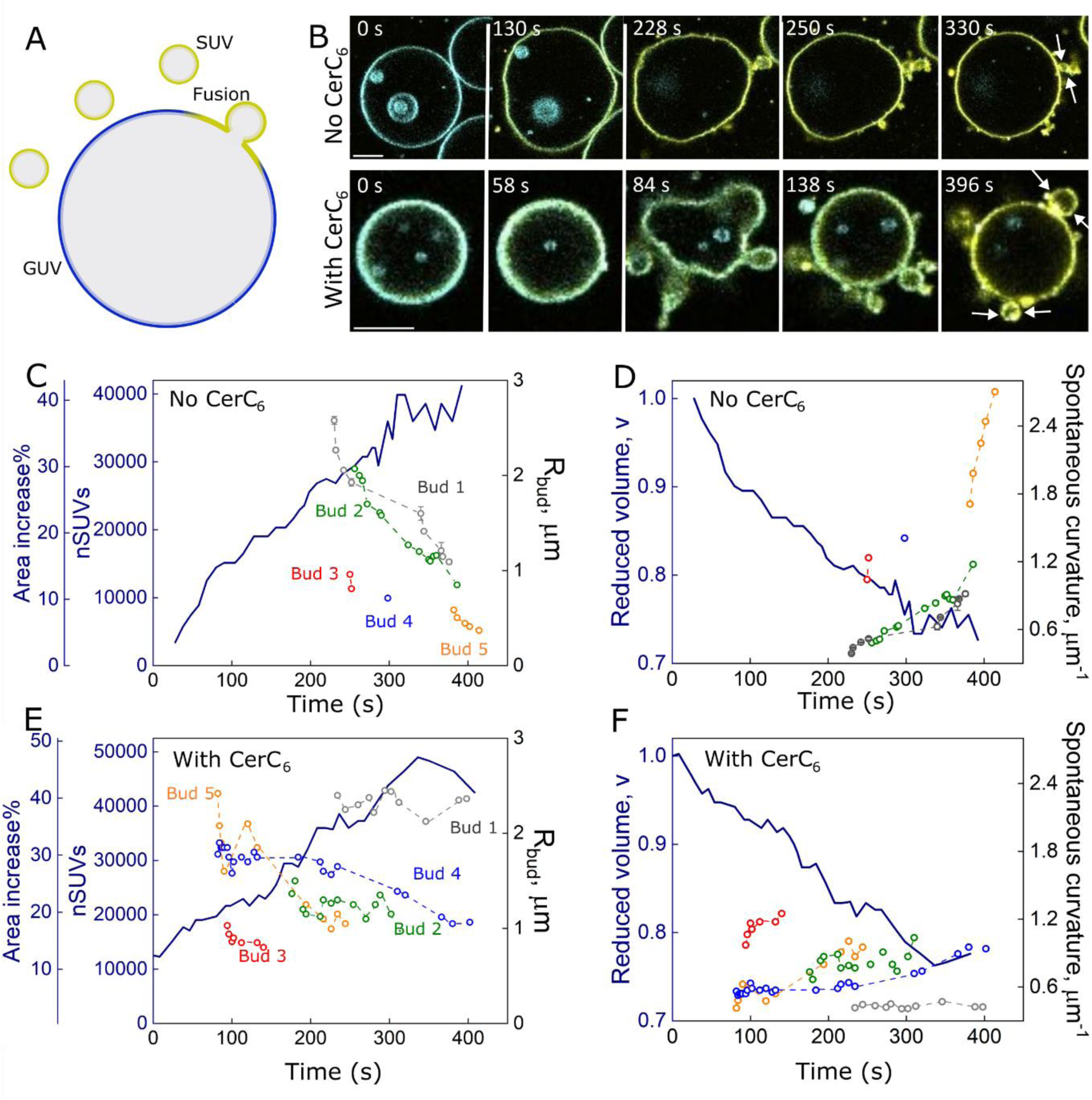
Real-time SUV-GUV fusions and assessment of mechanical parameters. A, sketch of the fusion assay with membrane-labelled SUVs (yellow) fusing with single blue-labelled GUVs. B, representative single GUVs (labelled with 0.5 mol% Atto 390) fusion experiments for GUVs devoid of (upper panel) or containing (lower panel) 10 mol% CerC_6_. The SUVs (containing 2 mol% DOPE-Rh) were composed of DOTAP:DOPE (1:1 mol ratio). For fusion with GUVs containing CerC_6_, the SUVs also contained 10 mol% CerC_6_ (with the equivalent removal of DOPE). Time values correspond to the time from the onset of observation. Scale bars 10 μm. C and D, calculated nSUVs and percentual increase in GUV area as a result of fusion (solid line) for the GUVs shown in panel B. The measured bud radius R_bud_ are shown for 5 independent buds. E and F, calculated reduced volume (*v*, solid line) and spontaneous curvature (*m*) for individual buds on the vesicles shown in B.

Figure 2C shows the number of SUVs, nSUVs, for membranes devoid of CerC_6_. This shows nSUVs values up to ∼ 40,000 SUVs fusing to the GUV. The area increase of the GUVs due to the fusions was very significant, with up to a 40% area increase counting from the initial GUV area before fusion. The associated reduced volume *v* is shown in Figure 2D. For the example of the fusion displayed in Fig.2B, *v* was seen to decrease from *v* = 1.0 to 0.7 (solid line in Fig. 2D). Fusion also resulted in bud formation, whose size can be directly measured from the images, and from which *m* can be calculated. Figure 2C (colored symbols; right axis) shows the evolution of R_bud_ for several independent buds. From the data, it is apparent that R_bud_ decreased as more SUVs fused. Figure 2D (right axis) shows the calculated *m* for each of the buds, with *m* strongly rising with time, with values ranging from 0.4 to 2.7 μm^-1^.

CerC_6_-containing GUVs underwent a more dramatic shape change, with a sudden bud transition, see e.g. the example in Figure 2B and movie S2 and movie S3. The main difference between CerC_6_-containing and CerC_6_-devoid GUVs was the earlier occurrence of a budding event where more homogeneously sized buds were formed, whose radius was approximately constant over time. While the parameters (nSUV, increase in area, reduced volume) were comparable with and without CerC_6_ (cf. Figures 3C, 3E), indicating the CerC_6_ did not influence fusion efficiency, the curvature *m* = 0.8 ± 0.3 μm^-1^ (mean ± SD) was much lower with CerC_6_ compared to vesicles without CerC_6_ (where it strongly increased over time, with an average *m* = 1.5±0.7 μm^-1^; mean ± SD). These experiments show that SUVs fusion can result in a very significant increase in GUV area, and that lipid flip flop significantly relaxes the intrinsic membrane curvature.

**Figure 3.**
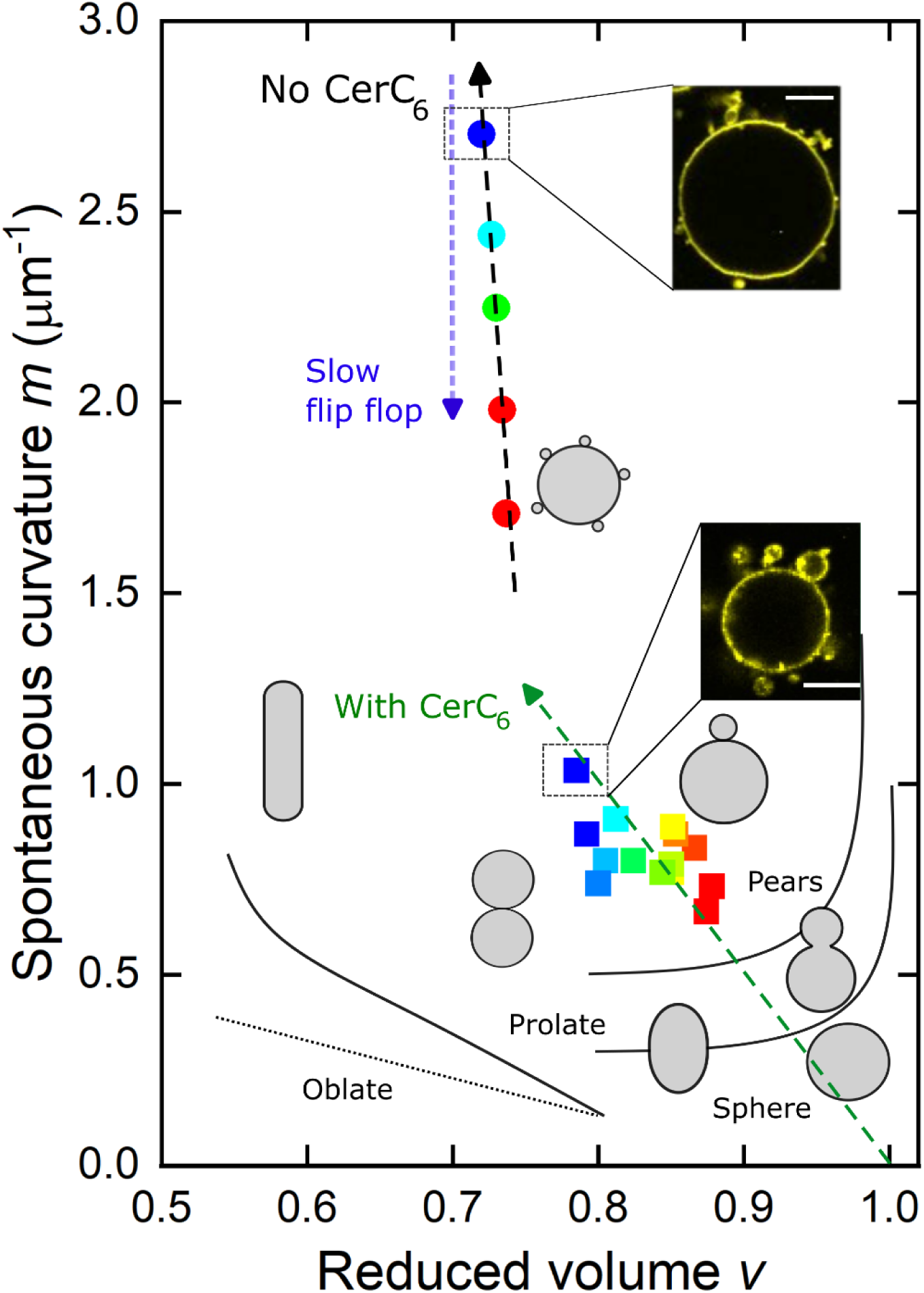
Tracking the shape trajectory of growing GUVs. Coloured data points represent the temporal evolution of GUVs devoid of (circles) and containing CerC_6_ (squares). Colors represent the time that evolves from red to blue over a timeslot of 382-415 s for the GUV without CerC_6_,, and 176-310 s for the GUV with CerC_6_. The green dashed arrow represents the trajectory for the CerC_6_-containing GUV, whereas the black arrow represents the trajectory of the vesicles without CerC_6_. The blue arrow represents the trajectory of delayed flip flop over longer timescales. The insets show representative GUVs observed for each of these conditions. The experimentally observed shapes match those that were predicted theoretically (grey shapes). Scale bars 10 μm.

We used the observed changes in *v* and *m* from the experiments in Figure 2 to locate the GUVs within the *(v*, *m)* shape phase diagram over the time course of the SUV fusion processes, see Figure 3. The GUVs started the growth process near the bottom-right location, with *v* = 1 and *m* ∼ 0 (i.e., a perfectly spherical shape and symmetric transbilayer lipid distribution), whereupon they underwent shape changes over time as SUVs fused to them. Notably, the quantitative data can only be shown after the budding transition since *m* is assessed from R_bud_. For the GUV devoid of CerC_6_, the buds appeared at *v* ∼ 0.74, *m* ∼ 1.7 μm^-1^, whereupon *m* further increased sharply to 2.5 while *v* only slightly decreased to 0.72. In contrast, the GUV containing CerC_6_ exhibited a much milder change in *m* as v decreased, with a budding transition at higher *v,* i.e., near *v* ∼ 0.9, *m* ∼ 0.6 μm^-1^. The experimental shapes that we observed in different locations of the diagram (cf. insets in Fig.3 as well as Figs.1-2) matched well with the expected GUV morphologies from previously published data^9^.

We measured the same behaviour shown in Figures 2 and 3 for several GUVs, although these experiments are experimentally challenging (e.g. since in most GUVs, the buds move in and out of focus, making it difficult to assess *m* for the vesicles where R_bud_ could clearly resolved). Many GUVs exhibited a comparable 50% area growth as further demonstrated in Figure S1A, with Figure S1B showing the shape trajectories. In conditions of lipid flip flop, the GUVs grew (i.e., *v* decreased) in a process where much milder increases in transbilayer asymmetries (*m*) developed. All in all, these results are the first to dynamically map the vesicle shape trajectories in a growth process as a function of time. They show that flip flop alleviates mechanical stresses in growing vesicles, generating more relaxed and more symmetrical distribution of sizes between the formed buds and the mother vesicles.

### Division of grown vesicles

As described above, the growing GUVs undergo a morphological budding transition, forming buds due to changes in *m* upon SUV fusion. To assess whether these changes in *m* are sufficient to generate bud *division*, we first performed Fluorescence Recovery after Photobleaching (FRAP) to probe the neck connectivity of buds to the main GUV. We performed FRAP on a central region of GUVs that were encapsulating the aqueous probe sulforhodamine B (SRB, ∼1 nm diameter) that was contained within the GUV and its buds (Figure 4A). If the lumens were connected through an open neck, the fluorescence of the bud would decrease as non-bleached SRB from the buds would partially replenish the GUV lumen (Figure 4A, mid). Figure 4B shows a representative GUV (containing CerC_6_) before and after SRB photobleaching (Movie S4 shows the complete sequence). Notably, the bud fluorescence remained constant for large times (>1 minute observation times), showing that no SRB transport occurred across the bud neck. From simple SRB diffusion, one would expect a very short (∼0.1 s) timescale for content exchange for a typical bud with a 5 μm diameter. This indicates that the bud neck was closed, as sketched in Figure 4A, right. We then reversed the FRAP experiment by performing FRAP of the bud, where both fluorophores in the membrane and the bud lumen were bleached. Figure 4C sketches the expected outcome for an open neck (left), a neck that was closed for bulk transport whereas membrane diffusion could still occur across the neck (middle), and a separate bud that had divided off but was still connected (right). Surprisingly, the experimental observations supported the middle scenario, as the majority of GUVs (5 out of 6 of events probed with GUVs containing CerC_6_) exhibited lipid exchange across the neck (fluorescence recovery within a few seconds) whereas none of these GUVs exhibited content exchange (Figure 4D and Movie S5). Importantly, the formed buds were mechanically identical to their parental GUVs (Figure S2). The results indicate that the bud neck is closed for diffusion of soluble cargos while its membrane remains connected to the parental GUV.

**Figure 4.**
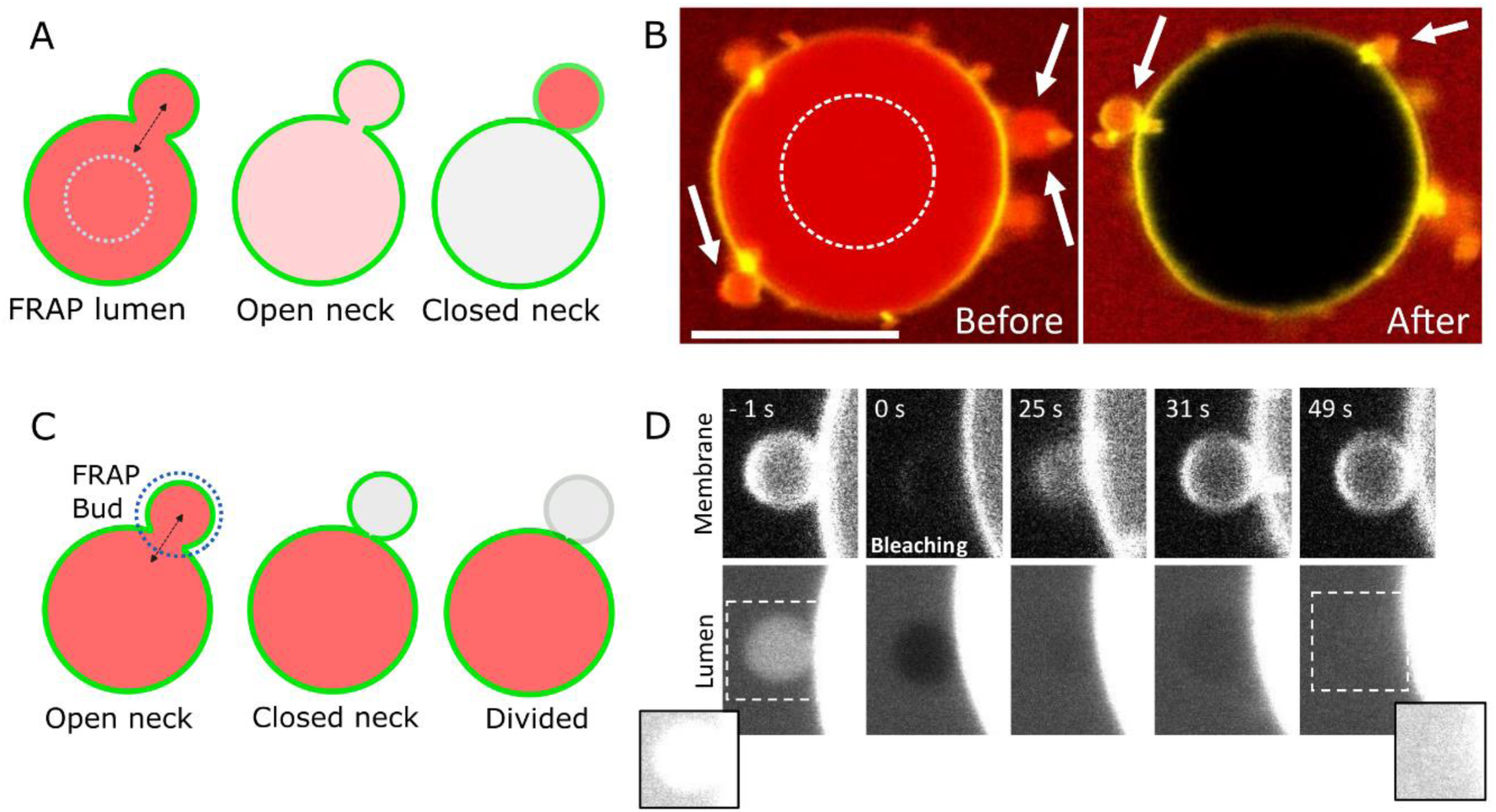
Transbilayer asymmetry generates buds with closed necks. A, Left: FRAP sketch (hashed circle) of GUV lumen filled with an aqueous probe (red) to check if bud neck is open. Middle: solution exchange dilutes SRB from the bud and partially recovers GUV lumen fluorescence. Right: a closed neck would not change intensity nor recover fluorescence in the GUV lumen. B, snapshots of a GUV before and after SRB FRAP within the hashed circle. The arrows point to some buds with stable lumen fluorescence. Scale bar 10 μm. C, FRAP of bud lumen and membrane to check for aqueous (red) and membrane (green) continuity. D, representative snapshots of the bud before (-1 s) and after bleaching of the membrane and lumen. The insets show brightness and contrast-enhanced images from the region within the hatched boxes. Scale bar 2 μm.

The growth of the synthetic GUVs showed the formation of highly curved buds that are only one step away from division. The spontaneous curvature *m* is predicted^33^ to generate a neck constriction force *F* of about *8πkm*, where^33^ *k* is the bending rigidity. From the curvature data in Figure 3, and a bending rigidity of ∼50 k_B_T for DOPC:DOPG membranes^34^, the force on the bud neck in the GUVs can thus be estimated to be ∼5 pN. For comparatively soft vesicles where curvature was induced by external protein binding^35^, division was reported to occur at neck forces of 26 pN (corresponding to *m* ≥ 6 μm^-1^), i.e., a value significantly higher than 5 pN. Therefore, the buds formed in our GUVs are not expected to spontaneously shed from the GUV, as indeed observed. For physical scission of the buds, an additional increment of *m*, and hence curvature force on the neck, is required.

To induce a further increase in membrane curvature *m*, we employed a recent approach based on rapid modifications of local lipid area upon light excitation of a photosensitive molecule (Figure 5A). The photosensitive molecule Chlorin e6 was incorporated in the outer leaflet of GUV membranes (Figure 5 A, ii), where it photooxidizes lipids upon light excitation, leading to an asymmetric increase in the outer leaflet area, which can induce membrane scission (Figure 5 A, iii and iv)^36^. Indeed, light activation of deflated GUVs in the presence of 100 μM external chlorin e6 led to GUV budding and scission in just a few seconds (Figure 5B and movie S6). After division, the separated bud slowly (tens of seconds) diffused away with no signs of lipid connection, as also assessed by the lack of any fluorescence recovery after photobleaching (Figure 5C and movie S7), confirming scission. Additional examples are shown in Figure S3, movie S8 and movieS9. Importantly, illumination in the presence of chlorin e6 did not cause membrane permeabilization (Figure S4), although, unlike previous reports, we found that chlorin e6 permeated through the GUV membrane, affecting the inner leaflet and (reversibly) quenching membrane fluorescence (see discussion in SI and Figure S5-S7).

**Figure 5.**
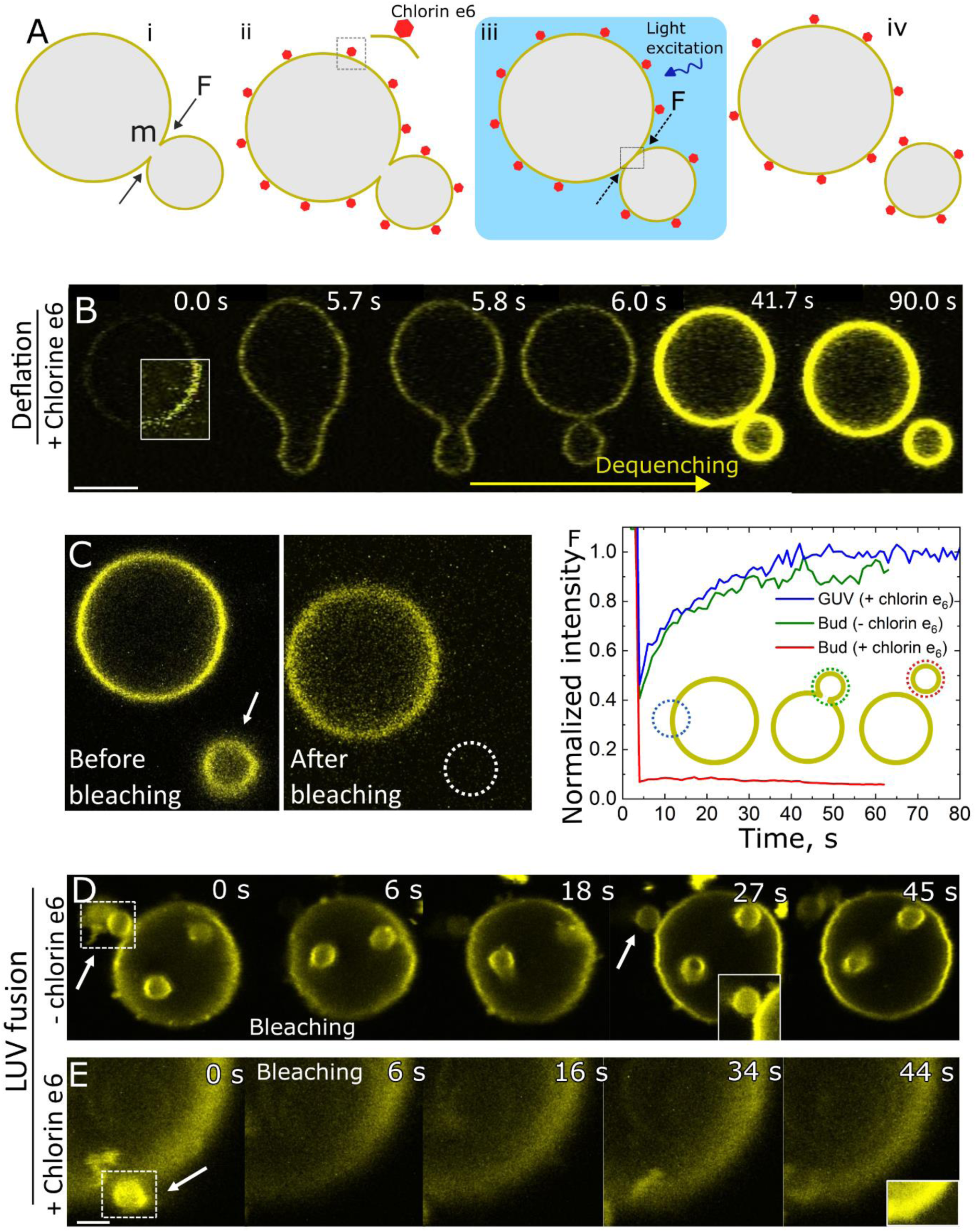
Division of grown GUVs. A, sketch of the division mechanism. (i) GUV buds formed after fusion gives rise to a constriction force F on the bud neck. (ii) Chlorin e6 mainly accumulates in the GUV outer leaflet. (iii) Photoactivation of chlorin oxidizes and increases lipid area, increasing the constriction force at the neck, thus (iv) severing the bud. B, deflated DOPC bathed in a 100 μM chlorin e6 solution divides upon illumination, which also de-quenches the fluorescent probe. Note that the GUV is initially highly quenched (as shown by the square with contrast-enhanced membrane). Scale bar 10 μm. C, image of the divided GUV shown in panel B, before and after photobleaching of the divided bud. Hatched circle: bud location after FRAP. D, photobleaching of a small chain of buds (arrow) formed after SUV fusion in the control sample without chlorin e6. The inset at t = 27 s shows a magnification of the recovered bud. E, FRAP of a divided bud. Note the lack of fluorescence recovery. The inset at t = 4 s show the highlighted area of bud location. F, evolution of fluorescence intensity after FRAP on a photobleached GUV membrane segment in the presence of chlorin e6 (blue line), on a control bud without chlorin (green line), and on a divided bud (red line). The inset sketches show the bleached segments with their respective colour codes.

We employed chlorin e6 to test whether it can be used to divide the buds from the GUVs that grew via fusion with SUVs. First, fusion with a relatively low SUV concentration (8 μM lipids) produced floppy GUVs prior to the budding transition. When illuminated with a strong 405 nm laser in the presence of chlorin e6, these GUVs divided. This process commonly resulted in multiple separated buds (Movie S10) and occasionally even proceeded symmetrically to form two equally sized GUVs (Movie S11). These results demonstrate that we can successfully divide GUVs that grew after fusion SUV fusion. However, with this low SUV concentration, the growth of the compartmental size was limited to only a few %.

We therefore tested if we are able to divide vesicles that formed as a result of more significant fusion and growth by incubating the GUVs with 30 μM SUVs (i.e. yielding a GUV area increase of 50% or more). For imaging, we mildly immobilized these GUVs in low-concentration agarose to minimize buds going out of focus^37^. Photobleaching of control buds (no chlorin e6) that formed after fusion revealed a fast and complete FRAP recovery (Figure 5D, green recovery in Figure 5F and Movie S12). Control FRAP in the presence of chlorin e6 but without 405 laser illumination also showed a full recovery (blue line in Figure 5F). In contrast, upon illumination with the 405 nm laser in the presence of 100 μM chlorin e6, we observed bud scission, as revealed by the complete lack of fluorescence recovery (Figure 5E, red line in Figure 5 F and Movie S13). In 16 out of 18 of such FRAP experiments, the buds successfully divided, i.e., the process occurred with a success rate of ∼90%, which is significantly higher than previous attempts with other approaches^35,38,39^.

The results show that GUVs that grew as a result of fusion with SUVs can be efficiently divided, by scission of symmetrical vesicles at lower *m*, or more asymmetrically upon bud scission. Note that these processes are governed entirely by physical mechanisms, showing that this can be achieved in synthetic cells without the need for a protein-based machinery as required in natural cells. This *in vitro* GUV growth and division are reminiscent of the process of cell division in L-form bacteria and possibly may recapitulate the division of primordial protocells that lacked divisome proteins. The findings provides guidelines for the division of man-made synthetic cells.

## Discussion

Cell division requires the duplication of cellular components, cellular growth, and the scission of the expanded cell into, typically symmetrically sized, daughter cells. In common eukaryotic cell lines^40^, cell growth mainly occurs in the G1 phase, with reported lipid biosynthesis lasting a few hours, while the cell cycle is reported to be much longer (∼1 day)^41^. The synthesized lipids are incorporated into the cytosolic leaflet of the ER and must move into the opposite luminal leaflet to allow for organelle growth before lipid redistribution to other organelles. In budding yeast^42^ and L-form bacteria^43^, the cell cycle occurs on a shorter timescale (∼1h)^44^ with the generation of highly curved cellular fragments (i.e. cell buds) that are initially connected to the parental cell, but eventually shed off, forming many daughter vesicles in progenitor L-forms^5,43,45^. In walled bacteria, which typically double in 0.3-1 hour, at least 5000 lipid molecules must flip every second across the membrane to meet the expanding demands of the growth ^14^. In the absence of an efficient flipping mechanism, synthesis would lead to a rapid accumulation of lipids in the cytosolic side of the bacterial membrane, generating inward buds that would disrupt faithful division.

Here, we recapitulated the morphological aspects of cell growth and division using a purely reconstituted system. Fusion of large amounts (tens of thousands) of SUVs to a single GUV, significantly increased the GUV area (∼50%) in only a few minutes. This initially led to visible GUV membrane fluctuations, which was followed by a budding transition due to a lipid number asymmetry build-up in the GUVs – generating GUVs that were covered with many small buds on their surface. We found that fast lipid flip flop alleviated curvature stresses during the growth process and led to the formation of more symmetrical GUVs with larger buds. The process was not specific to the particular flipping molecule as slow flip flop over long timescales led to similar relaxation in curvature stresses. Recent molecular dynamic simulations showed that the introduction of a lipid species that undergoes frequent flip flops enables the membrane to relax into a state where both leaflets are tensionless^46^. This is in line with our observations where membranes containing CerC_6_ were much less tense (i.e. wider buds) than those devoid of CerC_6_, as also illustrated in the shape trajectories. The formed buds had an identical composition compared to the mother GUV and developed closed necks, preventing solution exchange between the GUV and the bud lumens, a configuration that closely resembles the morphology observed during the late stages of budding yeast division^47,48^. We showed that an additional increase in transbilayer asymmetry (by actively increasing the area of outer leaflet lipids) led to full scission of the bud neck, severing the buds with a remarkably high success rate of >90%.

The phenomena described here are not specific to fusion of SUVs to GUVs or the particular CerC_6_ employed , but carry some general significance. Processes that are associated with transbilayer lipid asymmetries and curvature stress are ubiquitous in the cell. Given the high amounts of cholesterol in the plasma membrane (∼ 40%^18,49^), and considering that cholesterol exhibits fast flip flop in the order of ns^50^ (similar to CerC_6_; Ref.23), transbilayer asymmetries that are generated by fusion of small vesicles (which change surface area and curve and perforate the membrane^51^) or by direct lipid incorporation upon synthesis, would be nearly instantaneously buffered by cholesterol. Indeed, cholesterol has been recently shown to regulate the stability^52^ and shape of synthetic and living cells^19^, again suggesting a role of lipid flip flop as a regulatory mechanism of membrane shape^57^.

Summing up, we demonstrated that lipid flip-flop relaxes asymmetries in lipid number across the bilayer. Rapid flip-flop facilitates stress relaxation in rapidly growing synthetic cells, generating a more symmetrical distribution of progeny cells. The buds formed upon fusion of small vesicles primed the membrane for division. Our findings indicate a viable way forward for growth and division of synthetic cells as well as suggest that, under favourable conditions, primordial protocells may have been able to grow and divide without any complex protein machineries.

## Materials & Methods

All materials and chemicals were used as obtained. Glucose, sucrose and the fluorescent dye sulforhodamine B (SRB) were purchased from Sigma-Aldrich (St. Louis, MO, USA). The lipids 1,2- dioleoyl-3-trimethylammonium propane (DOTAP), 1,2-dioleoyl-sn-glycero-3- phosphoethanolamine (DOPE), 1,2-dioleoyl-sn-glycero-3-phosphocholine (DOPC), 1,2-dioleoyl-sn-glycero-3-phospho-(1’-rac-glycerol) (sodium salt) (DOPG), 1-palmitoyl-2-oleoyl-sn-glycero-3- phosphocholine (POPC), 1-palmitoyl-2-oleoyl-sn-glycero-3-phospho-L-serine (sodium salt) (POPS), N-hexanoyl-D-erythro-sphingosine (CerC_6_), and the fluorescent probes 1,2-dioleoyl-sn-glycero-3-phosphoethanolamine-N-(lissamine rhodamine B sulfonyl) (ammonium salt) DOPE-Rh and 1,2-dioleoyl-sn-glycero-3-phosphoethanolamine-N-(7-nitro-2-1,3-benzoxadiazol-4-yl) (ammonium salt) DOPE-NBD were purchased from Avanti Polar Lipids (Alabaster, AL). Chlorin e_6_ was purchased from Cayman Chemicals (Ann Arbor, MI). The probe DOPE-Atto 390 was purchased from Atto-Tec (Siegen, Germany). Bodipy C_12_ was kindly donated by Klaus Suhling (King’s College London).

Small unilamellar vesicles (SUVs) were prepared by the hydration method followed by extrusion^53^. In short, a 2 mM lipid mixture containing the respective lipids were dissolved in chloroform under N_2_ atmosphere and stored at -20C before use. For SUV preparation, an equivalent amount of the lipid solution was placed in a glass vial and chloroform was evaporated under a N2 stream. If not stated otherwise, the SUVs were made of DOTAP:DOPE (50:48 molar ratio), or DOTAP:DOPE:CerC_6_ (50:48:10 molar ratio) and labelled with 2 mol% DOPE-Rh. Chloroform residues were further removed under vacuum for ∼ 2h, forming a dry lipid film. The lipid film was hydrated with a 200 mM sucrose solution at room temperature, vigorously vortexed for 30-60s and extruded 11 times through a 100 nm polycarbonate filter with the help of a manual extruder (Avanti Polar Lipids, Alabaster, AL). The typical stock lipid concentration was also 2 mM, and when required, the solution was diluted in the same sucrose solution. The formed SUVs were stored at 4C and used withing 2-3 weeks.

The GUVs were prepared by the electroformation method^54^ with minor modifications. Briefly, 10- 20 μl volume of a 2 mM lipid solution in chloroform was spread on the surface of two indium titanium oxide slides and evaporated under a stream of N_2_. The slides were sandwiched with the help of a Teflon spacer, creating a volume of ∼ 1.8 ml. The lipid films were then hydrated with a 200 mM sucrose solution and connected to a function generator to apply a 5 mV nominal Vpp voltage and an AC sinusoidal current of 10 Hz for 1-2h, after which time the GUVs were harvested and ready for use. The GUVs were made of (i) DOPC:DOPG (50:50 molar ratio) or DOPC:DOPG:CerC6 (40:50:10 molar ratio) and labelled with 0.2 mol% DOPE-Atto 390 or Bodipy C_12_. For SRB encapsulation in the GUVs, a sucrose solution containing 50 μM SRB was used for hydration of the lipid film.

The SUVs and GUVs were incubated (i) in bulk for end-point measurements, or (ii) in real-time under direct microscopy observation. In condition (i), 50 μl of the GUVs were incubated with increasing SUV concentrations [SUV] in a total 100 ml solution for 10-15 minutes. The samples were then harvested and placed on the microscope for immediate observation. For FRAP experiments on the GUV buds (see below), the incubated sample was mixed with agarose (prepared in 200 mM glucose) at a final 0.1 wt% agarose concentration to weakly immobilize the GUVs and reduce out-of-focus bud motion^37^. In approach (ii), 50 μl GUVs diluted in 200 mM glucose were added to the observation chamber and placed on the microscope for observation for a final 100 μl. After 5-10 minutes, the GUVs sank to the bottom of the chamber due to a density gradient. Then, a concentrated (2 mM lipid concentration) SUV solution (typically 2-10 μl) was added to the chamber and the samples were imaged immediately to follow the fusion process in real-time^55^. For the division experiments, the pre-incubated SUVs and GUVs were incubated with 100 μM chlorin e6 for 10 minutes and then the samples were mounted on the microscope. Division occurred upon 100% excitation with a 405 nm laser.

Confocal imaging was performed on a Nikon Eclipse Ti confocal microscope using NIS-Elements AR Nikon-software. Samples were imaged through a 60x water-immersion objective (1.27 NA). Images were acquired with a Galvano High Resonant Scanner at 512x512 pixels in a sequential mode to minimize bleed-through. DOPE-Atto 390 was imaged with a 405 nm laser line and its emission detected with a 425-475 nm band pass filter. Bodipy C_12_ was excited with a 488 nm laser line and its emission detected with a 500-550 nm band pass filter. DOPE-Rh was excited with a 561 nm laser line and its emission detected with a 570-620 band pass filter. We calculated the number of SUVs (nSUVs) the fused with the GUVs using a previously reported approach based on lipid mixing^29^, with

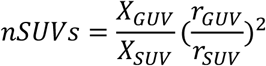

where X_GUV_ is the fraction of DOPE-Rh measured in the GUV after fusion, X_SUV_ is the fraction of DOPE-Rh in the SUVs (2 mol%), and r_GUV_ and r_SUV_ are the GUV and SUV radii, respectively. The data was pre-calibrated with an intensity response curve as a function of X_GUV_ in control GUVs (Figure S8) and the experiments were performed under identical imaging conditions.

Fluorescence Recovery After Photobleaching was performed in the FRAP mode of the AR Nikon- software, with 1-3 pre-bleaching images for a reference measurement at low laser power, followed by strong (100% laser power) illumination for 1 iteration (1 s/iteration), after which the samples were again imaged at low laser power for 30-60s. FRAP was performed on SRB encapsulated in the GUVs using the same imaging settings as for DOPE-Rh imaging, or on Bodipy C_12_ in the membrane buds using the same settings as above. The samples were analyzed using Fiji software for image intensity analysis, shape and GUV and bud dimensions. Roundness was measured after image thresholding and segmentation, from which the values are automatically obtained.

Fluorescence lifetime imaging microscopy (FLIM) and time-resolved fluorescence anisotropy were performed on a Microtime200 microscope (PicoQuant, Germany) using the Symphotime64 software as described previously^56^. Bodipy C_12_ was excited with a 485 nm laser operated at a repetition frequency of 40 MHz and 10 µW power as measured at the imaging plane. The emission light was passed through a 50 µm pinhole, split by polarization, and filtered by a 525/50 band pass filters (Chroma, USA) before being focused on single-photon avalanche-diode detector (PD5CTC and PD1CTC, Micro Photon Devices, Italy). GUVs were first localized using widefield imaging and then imaged in confocal mode using an objective piezo monodirection scanner, typical 200x200 px size (0.4 μm/pixel) and 0.6 ms pixel dwell times. The data was analyzed in the PAM software package using MATLAB^57^. The bright rim of individual GUVs was selected using a combination of intensity thresholding and manual selection. The intensity decays of the parallel and perpendicular detection channels were globally fitted using a model combining a bi-exponential fluorescence intensity decay with a single-exponential anisotropy decay using the approach of Schaffer et al.^58^

## Acknowledgements

We thank Ewa Pałczyńska for her help in performing preliminary division experiments, Anders Barth for assistance with spectroscopic experiments, Jacob Kerssemakers for discussions on data analysis, Klaus Suhling (King’s College London) for kindly donating the probe Bodipy C_12_, and Charu Charma, Bert van Herck, and Gijsje Koenderink for discussions. We acknowledge funding by BaSyC and the EVOLF SUMMIT project 1.004 that is financed by the Dutch Research Council (NWO).

