## Supplementary information for "Lipid flip flop regulates the shape of growing and dividing synthetic cells"

to the paper

### SI text

### Effects of Chlorin e6 on fluorescence quenching and membrane properties

We noticed that in the presence of chlorin e6, the GUVs were very dim compared to control samples, while strong illumination with 405 nm light caused a brightening of the GUVs, cf. Figure 5B in the main text. This effect was observed both for GUVs labelled with DOPE-Rh or Bodipy $C_{12}$ (Figure S4 A), although it had not been observed before[1]. Illumination partially increased brightness to control values, and brightening kinetics (Figure S4 B), and extent (Figure S4 C) depended on the probe used.

To gain insights into this brightening effect, we performed FLIM and time-resolved fluorescence anisotropy experiments (Figures S5 A and Figure S5 B). Chlorin e6 reduced the fluorescence lifetime of Bodipy $C_{12}$ from 2.0±0.1 ns to 0.6±0.2 ns (means and s.d.), an effect that is compatible with collisional quenching[2]. Since Bodipy $C_{12}$ is viscosity sensitive, its lower lifetime may result from fluorescence quenching, or membrane fluidization[3,4] (or their combination). If fluidization is the cause, fluorescence anisotropy should decrease due to increased probe rotation in more fluid membranes. Conversely, if it is due to quenching, then the anisotropy should increase because the probe will have less time to rotate (and hence lose anisotropy) while in the excited state. Figure S5 C shows that the rotational correlation time increased from 1.4±0.1 ns to 9.0±4.2 ns, indicating the second scenario.

Interestingly, the effects on lifetime and anisotropy was observed not only for the membrane directly exposed to chlorin e6, but also to membranes contained inside the GUVs (Figure S5 D), although to a lower extent, suggesting that chlorin e6 is able to traverse the membrane and therefore also acts on inner GUVs. In fact, vesicles inside vesicles also divided upon illumination (Movie S10 and Figure S6).

### Buds formed after fusion are mechanically identical to their parental GUVs

Fluorescence lifetime imaging microscopy (FLIM) on Bodipy $C_{12}$[3,4], a membrane molecular rotor that changes its fluorescence intensity and lifetime as a function of the local membrane viscosity, shows that intensity and lifetime were indistinguishable in the bud compared to the parental GUV (Figure 2A). This shows that they exhibit identical properties (e.g., viscosity, composition). The lifetime in the main GUV membrane was 2.07±0.01 ns while it was 2.10±0.01 ns (means and s.d.) in the bud (Figure 2B). Using previously reported calibration data[4], these values correspond to a viscosity of about 290 cP, comparable to values from liquid disordered membranes[5].

**SI Figures**

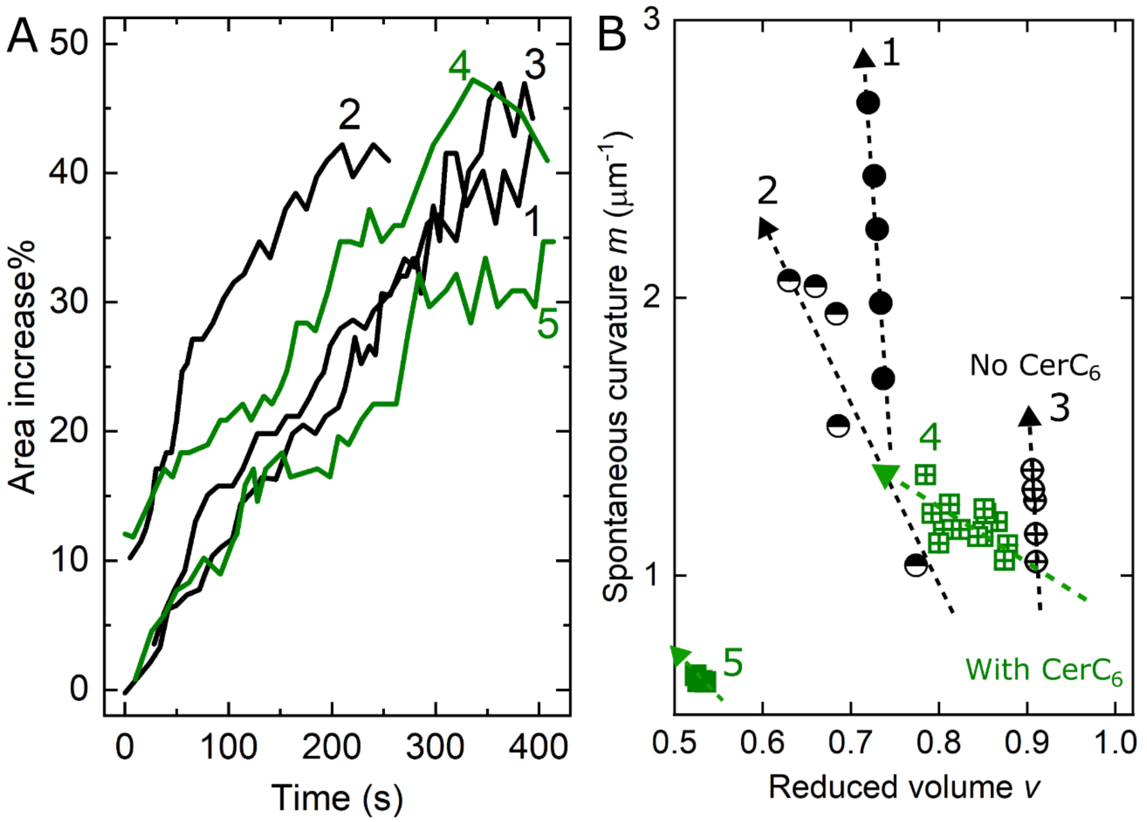


**Figure S1.** Tracking the shape trajectory of several growing GUVs. A. Measured GUV area increase from SUV fusion for five independent GUVs. Black and green represent vesicles devoid of and containing $CerC_6$, respectively. The observed growth for all GUVs was comparable and up to values of 35-45%. B. Evolution of the shape trajectory (arrows) for the growing GUVs shown in A. Note that $CerC_6$-containing GUVs gain area (i.e. reduced volume *v*) via SUV fusion with a much more moderate increase in spontaneous curvature *m* (GUVs 4 and 5), an indication of much milder tension compared to their GUV counterparts devoid of $CerC_6$ (GUVs 1-3).

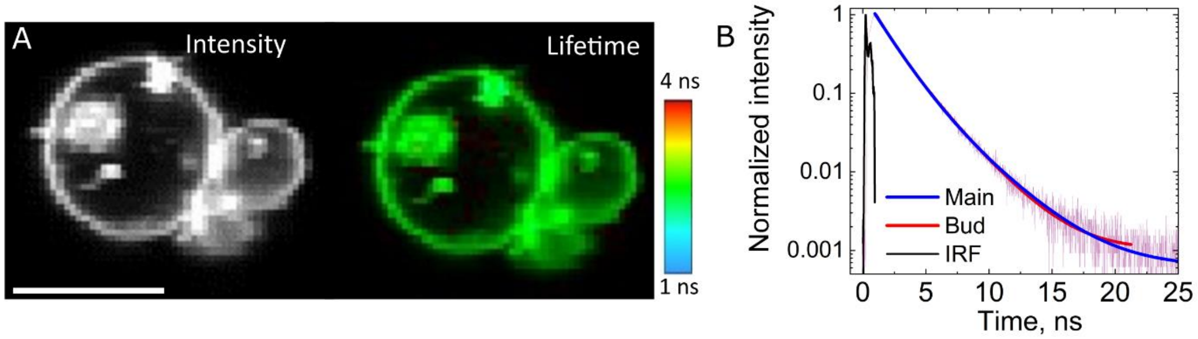


**Figure S2.** The formed buds are mechanically identical to their parental GUVs. A, fluorescence intensity and lifetime for a budded GUV. Scale bar: 10 μm. B, fluorescence lifetime measured on the main GUV membrane and on the bud. The decays, fits and calculated instrument response function (IRF) are shown.

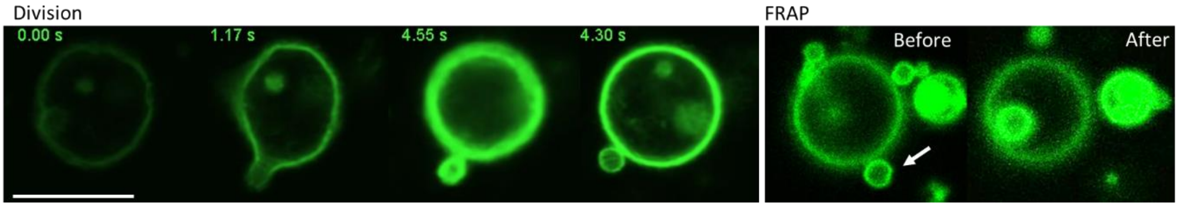


**Figure S3.** Division of a deflated DOPC GUV (labelled with 0.2 mol% Bodipy $C_{12}$) bathed in a 100 μM chlorin e6 upon strong 405 nm laser illumination (left panel). Note that the GUV becomes brighter upon illumination due to Bodipy $C_{12}$ dequenching. The right panel shows FRAP of the divided smaller vesicle before and after (1 minute) photobleaching. Scale bar 10 μm.

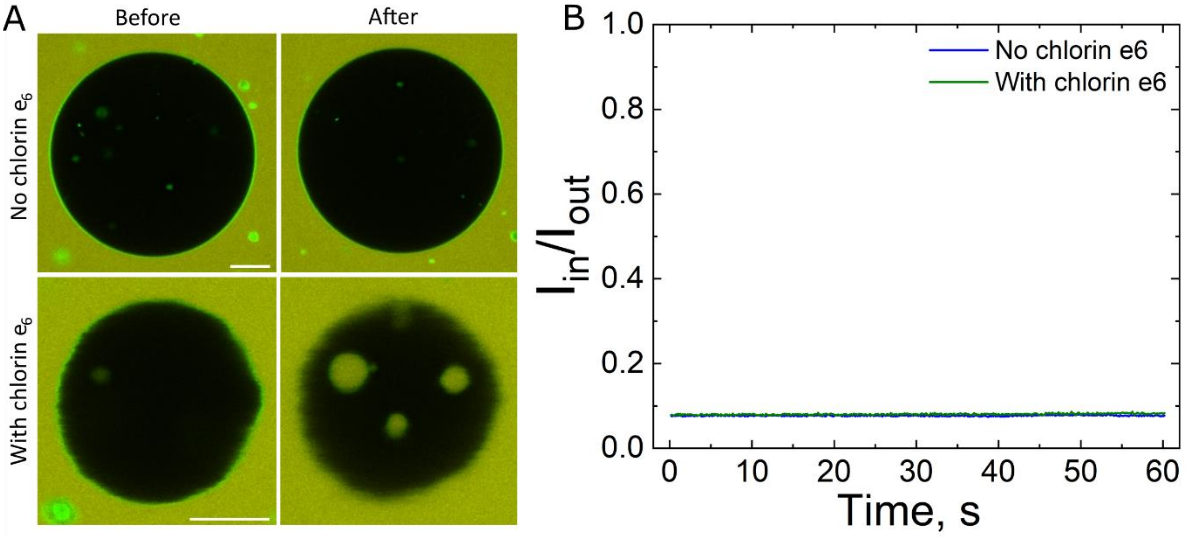


**Figure S4.** Illumination of DOPC:DOPG:CerC6 (4:5:1, mol ratio, labelled with 0.2 mol% Bodipy $C_{12}$) with 405 nm laser light in the presence of 20 μM SRB in the external medium. A, Images of representative GUVs before (left) and after (after) illumination for control samples (no chlorin e6, top) or incubated for 10-15 minutes with 100 mM chlorin e6 (bottom). Illumination of GUVs in the presence of chlorin e6 results in morphological changes, as the GUV buds inwards (encapsulating the external solution in doing so). Scale bars 10 μm. B, SRB intensity ratio inside ($I_{IN}$) and outside ($I_{OUT}$) for the two vesicles shown in A. These data indicate that SRB does not enter the GUV lumen, i.e., the membrane does not become permeable.

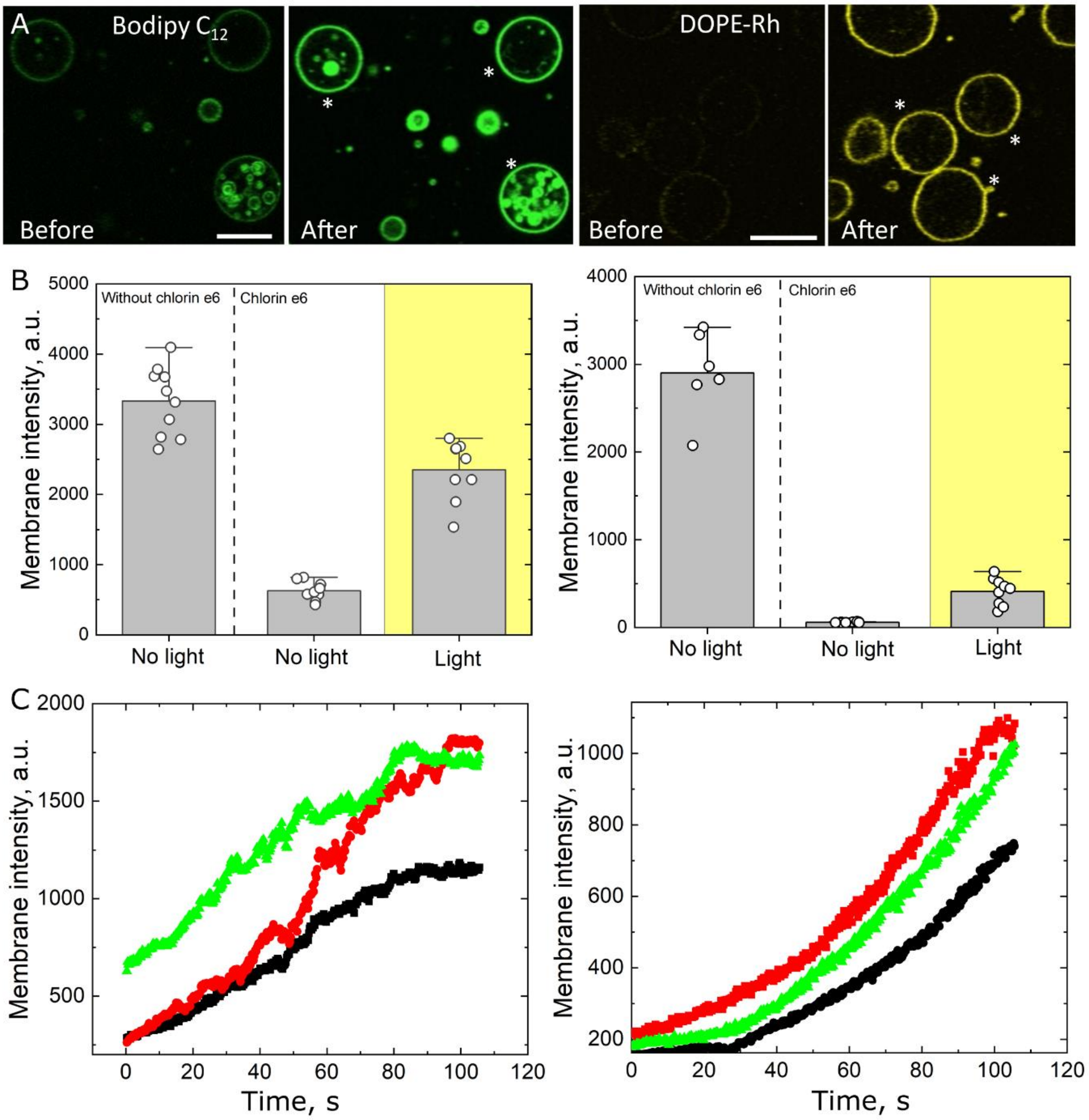


**Figure S5.** Chlorin e6 quenches GUV fluorescence and this effect is partially reversed by strong illumination. A. Representative images of DOPC:DOPG:CerC6 (4:5:1, mol ratio) GUVs labelled with Bodipy $C_{12}$ (0.2 mol%) (left, green) or 0.2 mol% DOPE-Rh (right, yellow) before and upon 1 min illumination with 405 nm laser light. Scale bar: 20 μm. B. Recovery of fluorescence intensity for the three vesicles for each labelled sample marked with an asterisk. C. Mean membrane fluorescence intensity for a number of individual GUVs labelled with Bodipy C12 (left) or DOPE-Rh (right), for control sample without chlorin e6 (left), control sample incubated with chlorin e6 without light illumination, and for sample incubated with chlorin e6 and upon light illumination. Each circle represents the measurement on an individual GUV, and bars are s.d.

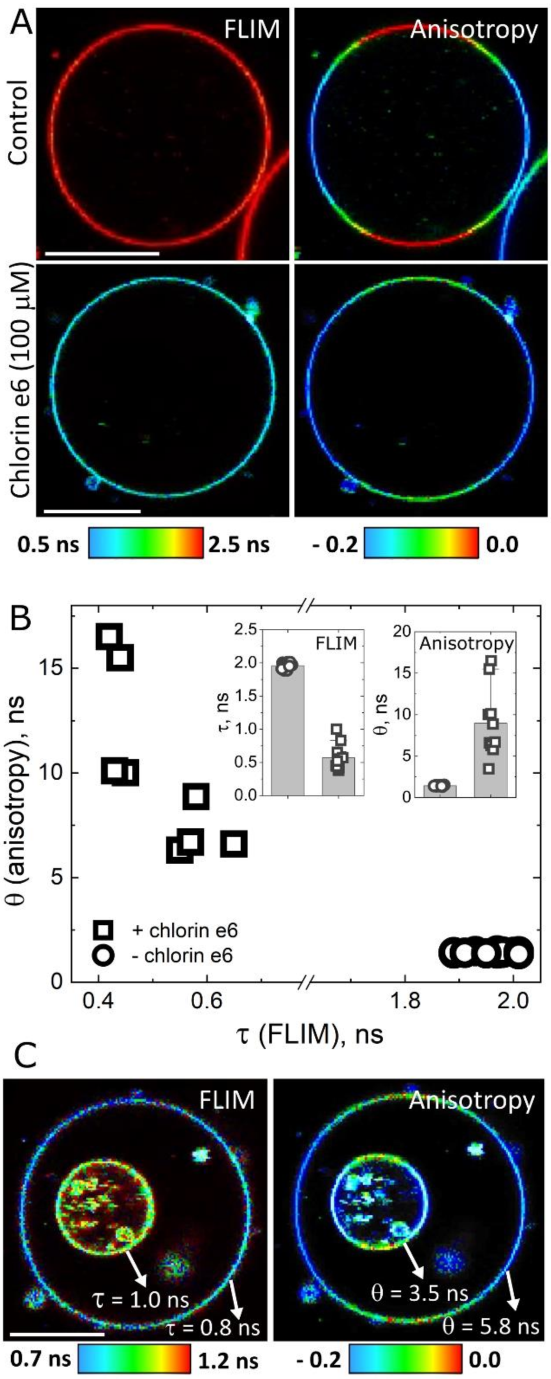


**Figure S6.** Chlorin e6 quenches fluorescence as assessed from FLIM and time-resolved fluorescence anisotropy. A. Representative FLIM and anisotropy images of DOPC:DOPG:CerC$_6$ (4:5:1, mol ratio) GUVs labelled with Bodipy C$_{12}$ (0.2 mol%) in the absence (control) and presence of 100 μM chlorin e6. Scale bar 20 μm. B. Measured rotational correlation time (θ) as a function of fluorescence lifetime (τ) for control (open circles) as well as for samples incubated with 100 μM chlorin e6. Inset: mean τ (left) and θ (right) values. C. Chlorin e6 permeates the GUV. The figure shows a representative GUV encapsulating a smaller GUV. The calculated values for the external and internal vesicles are also shown (arrows). Note that the fluorescence lifetime τ in the internal vesicle (1.0 ns) is significantly shorter than the average for several control vesicles as shown in Figure B (1.95±0.04 ns). For the outer membrane in this GUV, the measured τ = 0.8 ns. Similarly, the rotational correlation time θ is also significantly affected (3.5 ns) compared to the average values measured on control vesicles in Figure B (1.42±0.05 ns). For the outer membrane in this GUV, the measured θ = 5.8 ns

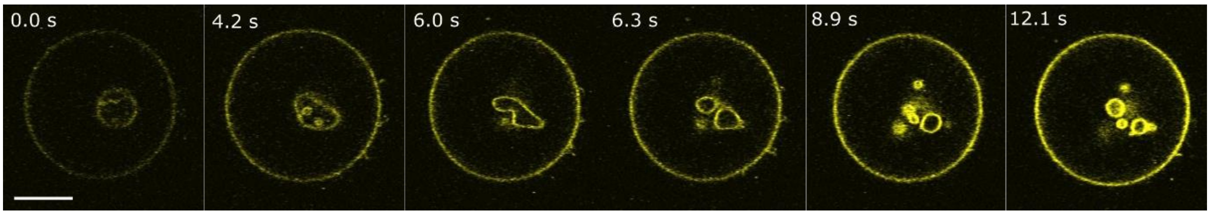


**Figure S7.** Chlorin e6 permeates the GUV membrane and divides internal vesicles upon illumination. Note GUV brightening upon 405 nm illumination. Scale bar 20 μm. The numbers at the top correspond to the time from the onset of illumination.

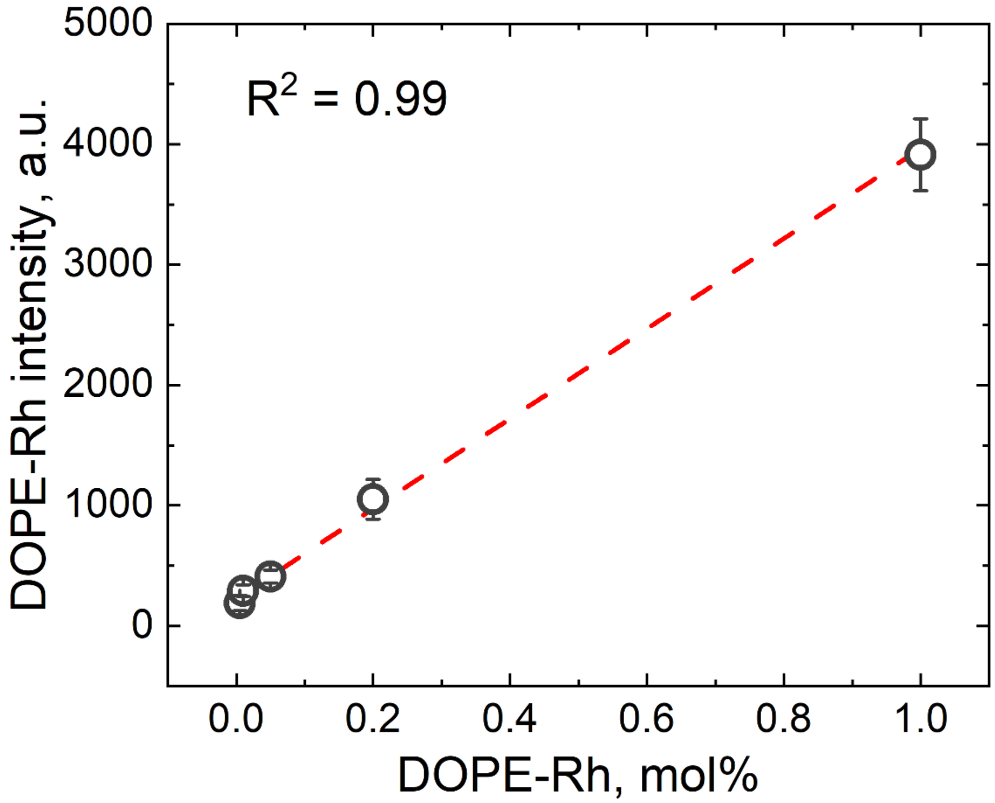


**Figure S8.** DOPE-Rh calibration curve. Measured DOPE-Rh intensity as a function of its mol% in the membrane of DOPC GUVs. 10-15 independent GUVs were measured for each condition. Means and s.d. are shown.

**Movie captions**

**Movie S1:** Growth of GUVs after fusion with many LUVs in the absence of flip flop. DOTAP:DOPE (5:5 mol ratio), labelled with 2 mol% DOPE-Rh (coloured yellow) were locally injected in a solution of DOPC:DOPG (5:5 mol ratio, labelled with 0.2 mol% DOPE-Atto 390, coloured blue). LUV fusion transfers its lipids to the GUVs, which grow and gain the yellow colour due to lipid mixing. The numbers represent the time from the onset of imaging. The diameter of the main GUV before fusion was 36 $\mu$m.

**Movie S2:** Growth of GUVs after fusion with many LUVs in the presence of flip flop. DOTAP:DOPE:$CerC_6$ (5:4:1 mol ratio), labelled with 2 mol% DOPE-Rh (coloured yellow) were locally injected in a solution of DOPC:DOPG:$CerC_6$ (4:5:1 mol ratio, labelled with 0.2 mol% DOPE-Atto 390, coloured blue). LUV fusion transfers its lipids to the GUVs, which grow and gain the yellow colour due to lipid mixing. Note the abrupt GUV budding. The numbers represent the time from the onset of imaging. The diameter of the main GUV before fusion was 20 $\mu$m.

**Movie S3:** Growth of GUVs after fusion with many LUVs in the presence of flip flop and "normal" budding. DOTAP:DOPE:$CerC_6$ (5:4:1 mol ratio), labelled with 2 mol% DOPE-Rh (coloured yellow) were locally injected in a solution of DOPC:DOPG:$CerC_6$ (4:5:1 mol ratio, labelled with 0.2 mol% DOPE-Atto 390, coloured blue). In this example, the GUV undergoes a less abrupt budding transition. The numbers represent the time from the onset of imaging. The diameter of the main GUV before fusion was 23 $\mu$m.

**Movie S4:** Photobleaching of the of the content marker SRB in the GUV lumen. Note that the SRB probe in the buds remain in the bud, demonstrating a lack of aqueous connectivity. The time represents the beginning of imaging. The GUVs made of DOPC:DOPG:$CerC_6$ (4:5:1 mol ratio, labelled with 0.2 mol% Bodipy $C_{12}$) and encapsulating 50 $\mu$M SRB were incubated with 15 mM LUVs (lipid concentration) made of DOTAP:DOPE (1:1 mol ratio, labelled with 2 mol% DOPE-Rh). GUV diameter was 14 $\mu$m.

**Movie S5:** Bud neck in growing GUVs formed after fusion is tight. Photobleaching of the of the lipid marker Bodipy $C_{12}$ in the membrane (left) and content marker SRB encapsulated (right) in the bud (arrow). Note recovery of the membrane marker but not of the content marker. The GUVs made of DOPC:DOPG:$CerC_6$ (4:5:1 mol ratio, labelled with 0.2 mol% Bodipy $C_{12}$) and encapsulating 50 $\mu$M SRB were incubated with 15 mM LUVs (lipid concentration) made of DOTAP:DOPE (1:1 mol ratio, labelled with 2 mol% DOPE-Rh). The time represents the beginning of imaging. Bud diameter was 2.7 $\mu$m.

**Movie S6:** Division of deflated GUVs upon light activation. DOPC GUV labelled with 0.5 mol% DOPE-Rh divides upon illumination of a strong 405 nm laser line in the presence of 100 $\mu$M chlorin e6. Note that the process occurs with simultaneous dye de-quenching. GUV diameter before division was 78 $\mu$m.

**Movie S7:** Photobleaching of one of the divided vesicles confirms division. Photobleaching of the smaller vesicle, which is physical separated and several $\mu$m distance from the other daughter vesicle. Note the absence of recovery. The GUVs are the product of the division shown in Movie S5.

**Movie S8:** Division of deflated GUVs upon light activation. DOPCDOPG:$CerC_6$ (4:5:1 mol ratio, labelled with 0.2 mol% Bodipy $C_{12}$) divides upon illumination of a strong 405 nm laser line in the presence of 100 $\mu$M chlorin e6. Note that the process occurs with simultaneous dye de-quenching. GUV diameter before division was 8 $\mu$m.

**Movie S9:** Photobleaching of one of the divided vesicles confirms division. Photobleaching of the smaller vesicle (arrow), does not exhibit recovery. The GUVs are the product of the division shown in Movie S5.

**Movie S10:** Division of internal vesicle when the GUV is bathed in a solution containing 100 $\mu$M chlorin e6.

**Movie S11:** Division of asymmetric GUVs that grew from LUV fusion. DOPC:DOPG:$CerC_6$ (4:5:1 mol ratio, labelled with 0.2 mol% DOPE-Rh) were incubated with 8 $\mu$M DOTAP:DOPE (5:5, labelled with 2 mol% DOPE-Atto 650, coloured red). The GUV is deflated due to an increase in area (rather than osmotic deflation). Strong 405 nm illumination in the presence of 100 $\mu$M chlorin e6 drives division through bud formation. Note that change from red colour, due to DOPE-Atto 650 photobleaching, and an appearance of a yellow colour, due to DOPE-Rh dequenching.

**Movie S12:** Division of symmetric GUVs that grew from LUV fusion. DOPC:DOPG:$CerC_6$ (4:5:1 mol ratio, labelled with 0.2 mol% DOPE-Rh) were incubated with 8 μM DOTAP:DOPE (5:5, labelled with 2 mol% DOPE-Atto 650, coloured red). The GUV is deflated due to an increase in area (rather than osmotic deflation). Strong 405 nm illumination in the presence of 100 μM chlorin e6 drives division to two equally sized GUVs. Note that change from red colour, due to DOPE-Atto 650 photobleaching, and an appearance of a yellow colour, due to DOPE-Rh dequenching.

**Movie S13:** Membrane buds formed upon LUV fusion are connected to the GUV. Control FRAP on buds formed on a POPC:POPS (5:5 mol ratio, labelled with 0.2 mol% NBD-PE). Note complete fluorescence recovery. The buds did not significantly diffuse away because the vesicles were immobilized in 0.2 wt% agarose. GUV diameter was 12 μm.

**Movie S14:** Membrane buds formed upon LUV fusion can be divided. FRAP on buds formed on a POPC:POPS (5:5 mol ratio, labelled with 0.2 mol% NBD-PE) GUV divide after membrane scission with a strong 405 nm laser excitation exhibit no fluorescence recovery, a sign of loss of membrane connectivity. The buds do not significantly diffuse away because the vesicles were immobilized in 0.2 wt% agarose. Note complete fluorescence recovery. Bud diameter was 1.3 μm.
